# Clusterin can mediate apoptosis-induced molecular mechanisms in immune thrombocytopenia

**DOI:** 10.1101/2023.09.26.559483

**Authors:** T. Stein, C. Bitsina, M. Schmugge, F. Franzoso

**Author notes:** Corresponding Author: Francesca Daniela Franzoso, Division of Hematology, Children’s Research Center, University Children’s Hospital Zurich, Zurich, Switzerland.

## Abstract

Abnormalities in the apoptosis pathway are possible risk factors for various autoimmune diseases including immune thrombocytopenia (ITP). ITP is an autoimmune bleeding disorder characterized by a low platelet count and mostly mild but in rare occasions life threatening bleeding symptoms. Platelets and megakaryocytes (MKs) may be seen as the major targets of the pathogenic immune responses in ITP. A mechanistic understanding of the ITP pathogenesis is still lacking. Our data indicate that mechanisms associated with impaired clusterin-mediated apoptosis might play a role in ITP platelet pathophysiology and platelet production by MKs.

We could demonstrate by apoptosis proteomic profiling significantly increased expression levels of some apoptotic genes such as clusterin (CLU), pro-caspase 3, catalase, TRAILR1/DR4, Bax, Bad and Bcl-2 compared to healthy controls in platelet-rich plasma (PRP) from 10 ITP patients. We could validate by both RT-qPCR and Western blotting that CLU, a stress-activated chaperone, is significantly increased in both newly diagnosed and chronic ITP. We used the human megakaryoblastic cell line MEG-01, treated for 4h with plasma from acute and chronic ITP patients and healthy controls. We performed chemical treatments in plasma treated MEG-01 by using pan-caspase inhibitors (Z-VAD-FMK), apoptosis inducer ABT-737, Rotenone and Rapamycin. We determined the expression at mRNA levels of apoptosis pathway regulatory genes Bax, caspase-3, -8, -9 as well as CLU, GRP78 and GRP94 by qRT-PCR. We could demonstrate significantly downregulation of mRNA expression levels of these apoptotic markers in ITP plasma treated and CLU siRNA transfected MEG-01 cells. Our results indicate a possible impairment of apoptosis pathway via upregulation of CLU and Bax in platelets and in their producers MKs that can lead to platelet destruction in ITP disease.

## Introduction

Immune thrombocytopenia (ITP) is an acquired autoimmune disease with a mostly benign course in children. ITP can occur in both children and adults and is more common in women than in men. It can present with severe mucocutaneous, in rare occasions even life-threatening bleeding symptoms ^1,2^. Classically, ITP is primarily caused by the production of autoantibodies against platelet cell surface molecules, such as the glycoprotein (GP) IIb/IIIa complex. In addition to autoantibody-mediated mechanisms, T cell-mediated destruction of platelets, antiplatelet antibodies and impaired megakaryopoiesis have been described as possible causes of the low platelet counts that hallmark this disorder. This highlights that the pathophysiology of ITP is for more complex than previously thought. Depending on its clinical course ITP is classified as a) an acute transient form termed as newly diagnosed (up to 3 months), b) persistent ITP (up to 12 months), and c) chronic form (more than 12 months) ^3^. In children, spontaneous remission resulting in a lower probability of disease recurrence and chronicity are common, whereas adolescents and adults tend to relapse and to develop chronic disease more often. The current goal of therapy is to increase platelet counts for the prevention of subsequent hemorrhages. As a first line therapy, intravenous immunoglobulins (IVIg), oral corticosteroids and anti-D immunoglobulins are currently used ^2^. However, there are many limitations to these treatments since many patients do not respond and/or develop persistent or chronic ITP. The factors that influence therapy response and clinical outcome are still unclear. Therefore, a better understanding of the underlying pathomechanisms could be a step towards the development of novel treatment strategies for ITP.

Platelets are produced by megakaryocytes (MK) in the bone marrow and, once released, they can undergo typical changes in their state of activation in the peripheral blood via platelet-glycoprotein-receptors ^4^. In platelets, as anucleated cells, apoptosis is mainly controlled by the intrinsic pathway through the interaction between the pro-apoptotic Bak and Bax and the anti-apoptotic prosurvival BCL-XL ^5^. The pro- and anti-apoptotic multidomain of the Bcl-2 family proteins regulates the entire platelet life span, whereas Bid and Bim are dispensable for Bax activation and mitochondrial apoptosis ^6^. However, we and others have not been able to demonstrate a role of receptor activation or of a “death ligand” initiating platelet apoptosis via the extrinsic apoptosis pathway ^7^. It remains unclear which or how proapoptotic signals induce apoptosis in platelets or even in their precursors, MKs ^8^. Previous studies also from our lab have reported the role of platelet apoptosis in the pathogenesis of ITP, demonstrating caspase-3, -8 and -9 activation in acute ITP ^9^ and increased phosphatidylserine (PS) exposure in chronic ITP ^5,9,10^. Further studies are needed to fully understand the mechanisms underlying these observations.

Clusterin (CLU) is a stress-activated chaperone that is highly expressed in Alzheimer’s disease and cancer and has been implicated in the pathogenesis of several protein aggregopathies and cancer entities ^11,12^. CLU can limit the severity of several autoimmune diseases such as autoimmune myocarditis, systemic lupus erythematosus and rheumatoid arthritis (reviewed in ^13^). CLU binds to histones expressed by late apoptotic cells and is involved in their clearance ^13^. Intracellular CLU localized to ER or mitochondria can directly bind to the pro-apoptotic Bax protein (Figure 2, ^14,15^). During stress conditions it can inhibit apoptosis by alleviating protein aggregation, and it can enhance Akt phosphorylation and trans-activation of NF-kB ^16,17^. Whether CLU is a pro-survival or a pro-apoptotic molecule remains unclear, it has most probably dual functions. CLU inhibitors are currently investigated in clinical trials in lung and prostate cancer ^16^. CLU can reduce the tumoricidal activity of histone deacetylase inhibitors tested for cancer treatment, by inhibiting apoptosis ^18^. CLU is expressed as different isoforms with distinct cellular or subcellular localizations in the brain ^19^. The CLU gene is organized into 11 exons (of which two are untranslated), spanning a region of 18,115 bp in humans. Transcription of CLU results in the production of three mRNA isoforms in humans and mice ^19^. The two primary isoforms, CLU1 and CLU2, have been first reported in the human brain, sharing exon 2-9 but differing in exon 1 and proximal promoters and presenting an increased expression in Alzheimer’s disease neuropathology ^11,15^. CLU was detected in alpha-granules of platelets and in bone marrow-derived megakaryocytes, most probably produced and packaged into the alpha-granules during megakaryocyte development ^20^. Previous reports showed that CLU can interact with Glucose-regulated protein (GRP78), a central regulator of unfolded protein response, playing an important role in cellular adaptation and survival under stress conditions, in particular in ER stress^21,22^.

We investigated if CLU together with GRP78, GRP94, Bax, p53, Caspase-3, -8 and -9 may be involved in platelet apoptosis in both newly diagnosed and chronic ITP. We further interrogate if similar mechanisms may be responsible for CLU-induced apoptosis in megakaryocytes treated with ITP plasma and if we could reverse these effects by RNA interference via siRNA transfections targeting CLU and GRP78.

## Materials and Methods

### Patient Characteristics

This study was approved by the local ethical committee (Cantonal Ethics Committee Zurich, Switzerland) and written informed consent was obtained from all healthy blood donors as well as from ITP patients, or their legal guardians. All patients were recruited at the Children’s Hospital Zurich, Switzerland.

All patients fulfilled the criteria for primary ITP (as defined in ^23,24^). In total, we recruited 12 acute ITP patients with median platelet count: 6.35 ± 5.79 ×10^9^/L; median age: 12.36 ± 5.20 years; gender ratio: 8 females (F) / 4 male (M)). For comparison, 15 chronic ITP patients (median platelet count: 77.60 ± 86.46; median age: 11.0 ± 4.14 years; gender ratio: 8F/7M) were studied. Chronic ITP patients were identified as patients with a persistent low platelet count (<100 × 10^9^/L), which lasted longer than one year. In addition, we included 11 healthy paediatric controls with a median platelet count of 263.5 ± 125.5 ×109/L and a median age of 10.5 ± 7.01 years (5F/6M) (as summarized in Table 1).

**Table 1.**
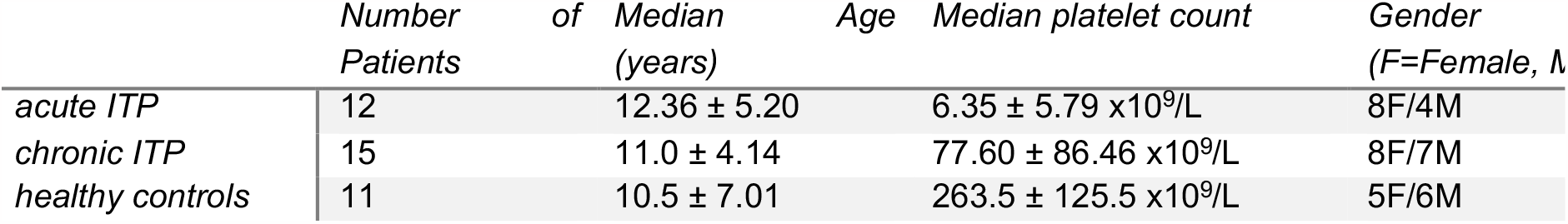
Summary of patient’s characteristics.

### Preparation of Platelet Rich Plasma (PRP)

First the citrated blood tubes were centrifuged at 150g for 20min at room temperature (RT). PRP was collected in a fresh 15ml falcon tube. Approximately 10ml of remaining blood was pooled in a 15ml falcon tube and frozen at -20°C for DNA. Then the PRP was centrifuged at 150g for 10 min at RT. The supernatant was carefully pipetted into a fresh tube, leaving the PRP pellet undisturbed. The tubes were labelled with date and sample ID and placed into the freezer at -80°C.

### PBMC Isolation

5ml of Ficoll (Sigma Aldrich) were prepared in a 15ml tube. The blood was diluted 1:1 or 1:2 with RPMI-1640 (Sigma Aldrich) medium and resuspended well. The diluted blood was carefully added on top of the Ficoll. The tubes were centrifuged for 20 min at 800xg with a fast start around 9 and a break at 1. After the centrifuging, the supernatant was carefully taken off and the PBMC monolayer was collected and placed in a new tube. The cells were once washed with PBS and centrifuged for 5min at 400xg at RT. 1-2 ml of RPMI medium was added to the PBMC, resuspended and the cells were counted.

### Proteomic apoptosis profiling

We used the proteome profiler human apoptosis array kit from R&D Systems (Mineapolis, MB, USA, Cat. ARY009) to assess apoptosis and apoptosis-related proteins according to manufactures instructions. In brief, PRP pellets were lysed in lysis buffer and their protein concentration was measured using Pierce BCA protein assay kit (Thermo Scientific, Cat. 23225) and a Tecan Infinite 200 plate reader (Tecan Group Ltd., Switzerland). The membranes were blocked with blocking solution and incubated overnight with the cell lysate (37.5μg). After the washing steps, the membranes were incubated with the biotinylated detection antibody cocktail and conjugated with streptavidin-horseradish peroxidase. To detect the signal intensity after adding chemiluminescent reagents we used the ChemiDoc Touch Imaging System (Biorad, CA, USA). For the analysis we used Biorad ImageLab 5.2.1. software.

### RNA Extraction and RT-qPCR Analysis

Total RNA was extracted from platelets and PBMCs using RNeasy mini kit from Qiagen, Germany as described in the manufacturer’s manual. Shortly, the cells were disrupted by adding QIAzol and homogenized by pipetting, vortexing and adding chlorophorm. Phase separation was achieved by centrifuging the samples for 15min at 12000xg at 4°C; the aqueous phase was collected and mixed with ethanol. For the RNA to bind, the sample was pipetted into a RNeasy Mini column and centrifuged at 8000xg for 15sec. The RNA was washed by adding RWT and RPE Buffer to the RNeasy Mini column and centrifuging the sample. To elute the sample, RNase-free water was added directly onto the RNeasy Mini column membrane. The RNA vials were put on ice and the nucleic acid concentration of the isolated RNA was determined using Nanodrop. The samples were then kept in -80°C. In a second step, the extracted RNA was reverse transcribed using the QuantiTect Reverse Transcription Kit from Qiagen as recommended by the manufacturer. RT-qPCR was performed in 96-well fast thermal cycling plates (Thermofisher Scientific, Switzerland) using PowerUp SYBR Green Master Mix (Thermofisher Scientific, Switzerland, cat. A25742) and primers with specific sequences as mentioned in Table 2.

**Table 2.**
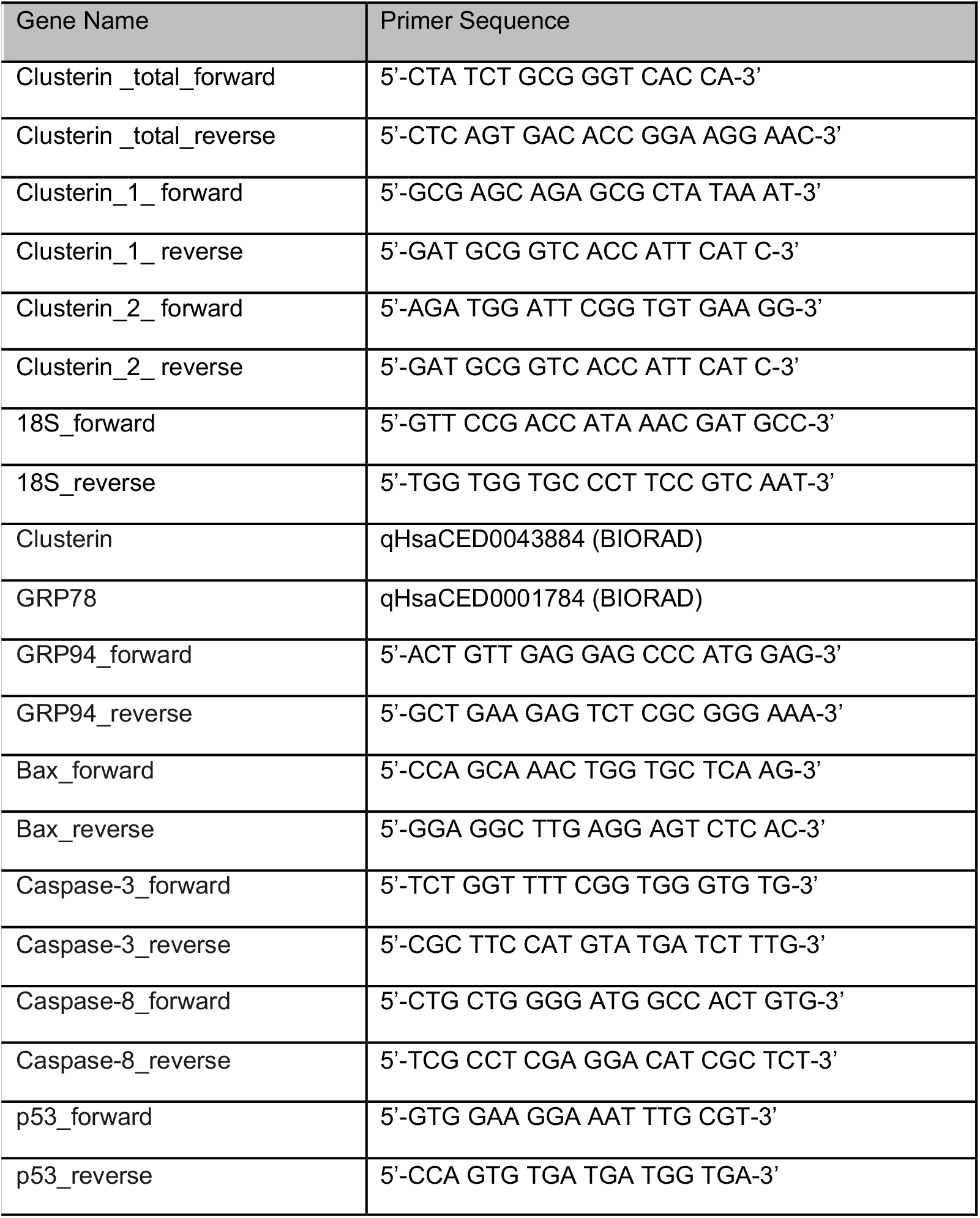
List of primers used for performed RT-qPCR.

### Western Blotting Analysis

Whole protein lysates were collected using lysis buffer for platelets (50 mM TrisHCl, 150 mM NaCl, 1 % Tween, 1mM EDTA, pH 7.4) and incubated for 2 min in ultrasounds. The lysates were gently rocked at 4 °C for 30 min and centrifuged at 14000 x g for 5 min and transferred into clean tubes. Bradford assay was performed for determining the protein concentration of each sample using Pierce BCA protein assay kit (Thermo Scientific, Pittsburgh, PA, USA, Ref.23225) and a Tecan Infinite 200 plate reader (Tecan Group Ltd., Switzerland). 12ul of protein sample were added to 3ul of 5x Lämmli buffer and denaturised for 5 min at 95°C. Gels (NuPAGE Novex Bis-Tris Gel 4-12%) were ran for 1-1.5 hours at 0.25A and 200V. We used Ponceau staining to mark the protein bands. After incubation for 1 hour with TBS (10ml TBS 20x, 190ml H20) and 5% milk at RT, the membrane was then incubated with relevant primary antibodies (1:5000) in 20ml TBS and 5% milk overnight at 4°C. After incubation, the membranes were rinsed 3 times in 15ml TBS-T (10ml TBS 20x, 2ml 10% Tween, 188ml H_2_O) for 5 min and then incubated with a secondary antibody (1:5000-1:10000) in 20ml TBS and 5% milk for 1 hour at RT. The membranes were rinsed 3 times in TBS-T for 5 min. The ECL substrate (Thermo Scientific, Cat. 32106) was added (1:1) and incubated for 1min. To detect the signal intensity after adding chemiluminescent reagents we used the ChemiDoc Touch Imaging System (Biorad, CA, USA). For the analysis we used Biorad ImageLab 5.2.1. software. Primary antibodies used are mentioned in Table 3 and secondary antibodies used are described in Table 4.

**Table 3.** Primary antibodies used for Western blotting.

| Name | Company | Host | Catalog number |
| --- | --- | --- | --- |
| Clusterin (A-9) | Santa Cruz<br>Biotechnology | Mouse IgG2a monoclonal | SC-166907 |
| Clusterin-α (A-11) | Santa Cruz<br>Biotechnology | Mouse IgG2a monoclonal | SC-166831 |

**Table 4.** Secondary antibodies used for Western blotting.

| Name | Company | Host | Catalog number |
| --- | --- | --- | --- |
| Beta-actin | Sigma | mouse | 066M4860V |
| Goat anti-mouse IgG (H+L) | Proteintec | Goat anti-mouse IgG (H+L) | SA00001-1 |

### Cell Cultures

The human megakaryoblast leukemic cell line MEG-01 (Sigma-Aldrich, ECACC 94012401) was maintained in RPMI 1640 Medium + GlutaMAX™-I (Gibco™ – Thermo Scientific) supplemented with 10 % of heat inactivated FBS (Sigma Aldrich, Cat. B9433). Cells were cultured in an incubator at 37 °C in a 5% CO_2_ humidified atmosphere.

### siRNA Transfection and plasma treatment

For siRNA transfections and chemical treatments, 40’000 cells per well were plated in a 24-wells plate. MEG-01 cells were incubated with siRNAs gene against Clusterin and negative control siRNA (Santa Cruz Biotechnology) for 48 hours according to manufacturer’s instructions. Four hours before harvesting, 40 μL of PRP plasma of a healthy control, ND ITP patient or chronic ITP patient were added to the cells. Cells were then collected and stored at -80 °C until further processing.

MEG-01 cells were incubated with pan-Caspase inhibitor Z-VAD-FMK (R&D Systems) up to a concentration of 10 μM, Rapamycin (BioGems) 10 μM or Rotenone (Sigma-Aldrich) 10 μM or with of ABT-737 (Active Biochem) 7. μM for two hours.

One hour before harvesting, 20 μL of plasma of a ND ITP patient, a chronic ITP patient or a healthy control, were added to the cells. Harvested cells were stored at -80 °C until further processing.

### Statistical Analysis

Our data was statistically analysed by using Prism 6.00 (GraphPad, Software CA, USA). Data are presented as mean +/-standard deviation of the mean from three independent experiments unless otherwise mentioned. Data was analysed using t-Tests or One-way ANOVA, followed by multiple comparisons tests to compare the means between the groups and p value < 0.05 was considered significant.

## Results

### Proteomic apoptosis profiling in ITP

We investigated the apoptotic proteomic profiling of platelets in both ND and chronic ITP patients compared to healthy controls (Figure 1). Several apoptotic genes such as Bcl-2, Procaspase-3, XIAP, TRAIL, CLU showed significant upregulation in ITP platelets when compared to healthy control platelets (Figure 1) suggesting that apoptotic mechanisms could be activated in the platelets from ITP patients compared to healthy controls.

**Figure 1.**
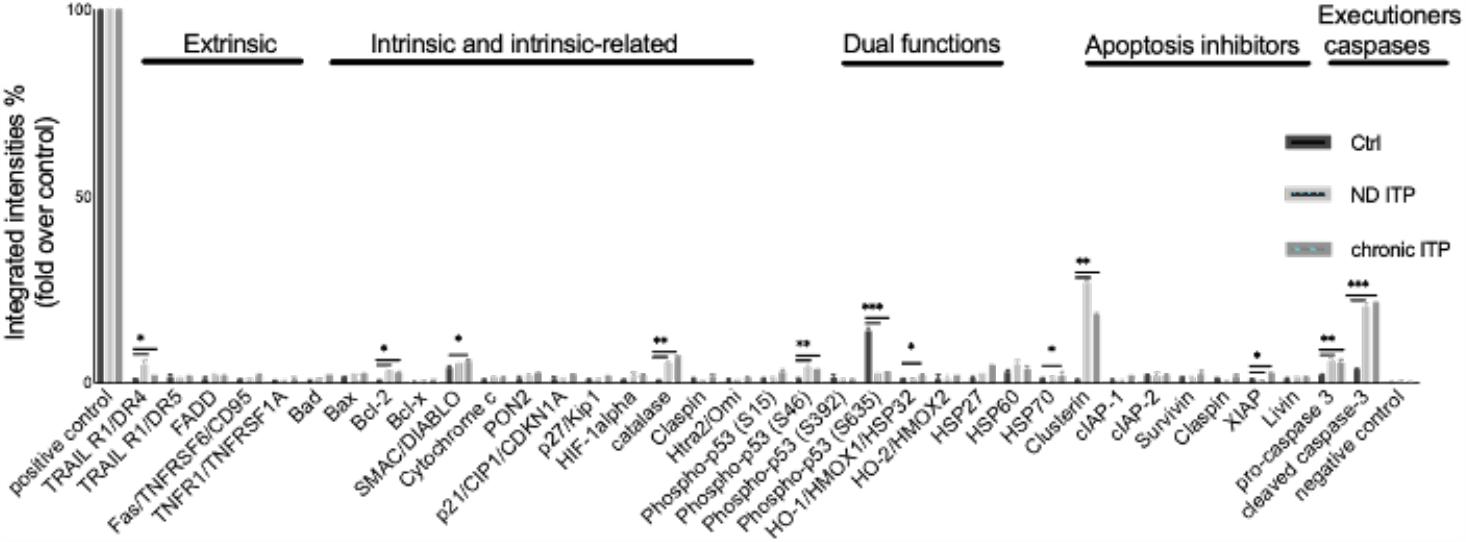
Apoplosis proteomic profiling in healthy controls, newly diagnosed (ND) and chronic ITP patients. Plalelets isolated from fresh PRP were analysed on a Human Apoptosis Antibody Array for the presence of apoptotic proteins (extrinsic, int,insic and intrinsic-related, dual functions. apoptosis inhibitors and executioners). The integrated densities of the dot blot assays were calculated according to Dot blot analysis (BioRad Image Lab 5.2.1.) and were normalia:ed to the positive control (representing 100%).

### CLU gene expression in ITP platelets and PBMCs

We validated the CLU expression results in platelets isolated from ND and chronic ITP patients at both mRNA and protein level by qRT-PCR and Western blot analysis (Figure 2A and B). Our results showed a significant increase in CLU as well as in the two isoforms CLU1 and CLU2 levels in both groups of ITP patients compared to healthy controls.

**Figure 2.**
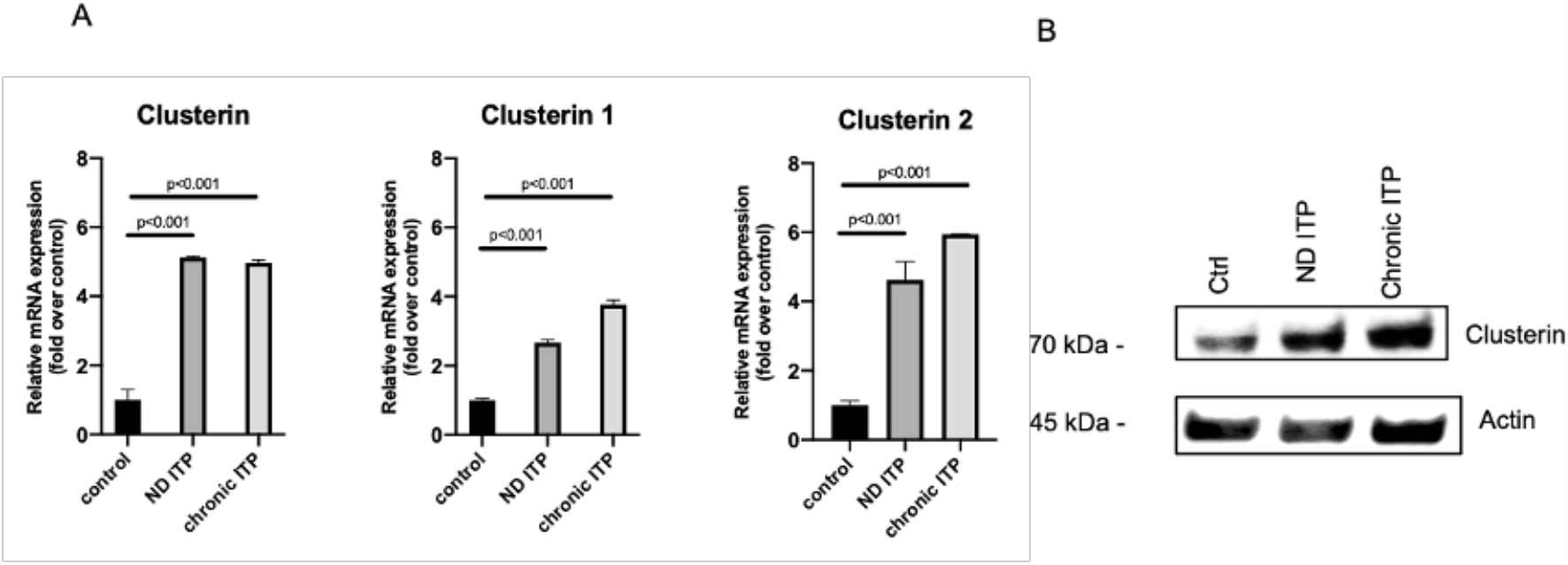
imRNA expression levels of Total Clusterin, Clusterin 1 and Clusterin 2 in platelets from our ITP patient groups. Total RNA from platelets was isolated and subjected to RT-qPCR with 18S rRNA as internal control. Relative mRNA levels of each sample normalized to healthy controls are shown as means (n=9) ± SD. Error bars show SD. Statistical analyses were performed using one-way ANOVA followed by multiple comparisons tests to compare the mean ranks between the groups. Means are significantly different with < 0.001 (highly significant) (B) Clusterin expression was assessed by Western blotting in healthy controls (lane1), newly diagnosed (ND) ITP (lane 2) and chronic ITP patients (lane 3). A representative immunoblotting image was selected (N=3).
.

Moreover, we also observed that CLU levels were significantly lower in peripheral blood mononuclear cells (PBMCs) from ND and chronic ITP patients compared to healthy controls, suggesting that the increased expression is specific to platelets (Figure 3).

**Figure 3.**
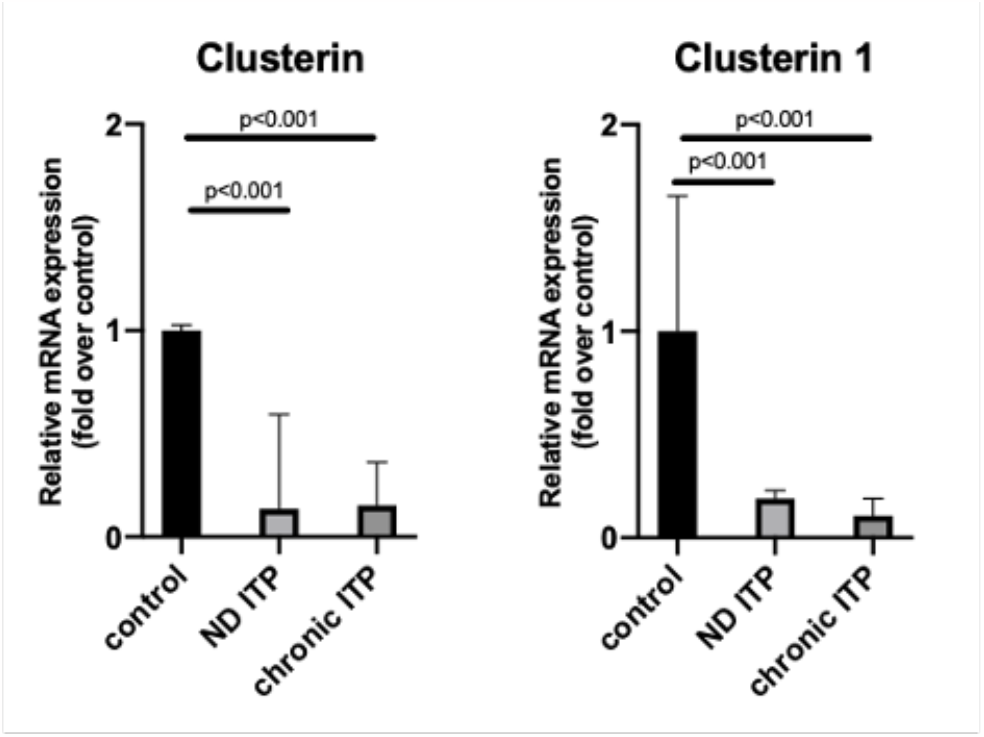
mRNA levels of Total Clusterin and Clusterin 1 in PBMCs from our ITP patient groups. Total RNA was isolated and subjected to RT-qPCR with 18S rRNA as internal control. Relative mRNA levels of each sample normalized to healthy controls are shown as means (n=2) ± SD. Error bars show SD. Statistical analyses were performed using one-way ANOVA followed by multiple comparisons tests to compare the mean ranks between the groups. Means are significantly different with p < 0.001 (highly significant).

### CLU activation mechanisms in ITP plasma treated megakaryocytes

We next proposed a further in-depth molecular analysis to determine which possible mechanisms are involved in the CLU-mediated apoptotic death of platelets in ITP disease and if MKs, as precursors of platelets, show similar changes as in platelets.

We used the megakaryoblastic cell line MEG-01 first treated with ITP or control plasma. In line with the proteomic data, we were able to confirm the upregulation of CLU, GRP-78, caspase-3 and p53 in the MEG-01 cell line treated with plasma derived from ND or chronic ITP patients compared to plasma isolated from blood of healthy individuals (Supplementary Figure 1).

To determine the effects of different apoptosis inducers on MEG-01 treated cells, we first used the mTOR complex I (mTORC1) inhibitor Rapamycin, known to induce apoptosis and autophagy in platelets ^25–27^. We observed a significant downregulation at mRNA level for CLU, Caspase-3, -8 and -9, p53 and Bax for the ITP plasma treated cells in comparison to control plasma or plasma untreated cells (Figure 4).

**Figure 4.**
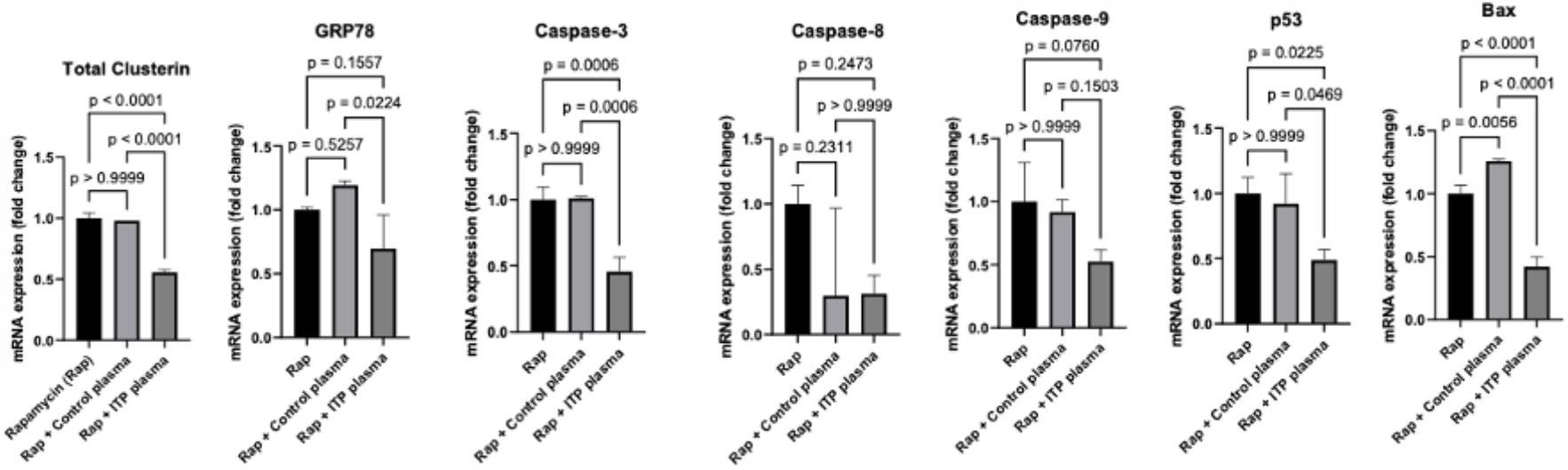
Effects of Rapamycin on apoptosis genes in MEG-01 cell line. Apoptosis genes were significantly downregulated in ITP plasma and rapamycin treatment. Relative mRNA levels of each sample normalized to control condition are shown as means ± SD. Error bars show SD. Statistical analyses were performed using one-way ANOVA followed by multiple comparisons tests to compare the mean ranks between the groups.

Secondly, we used Rotenone, a known apoptosis inducer ^28,29^ to compare changes that might occur in MEG-01 treated cells. Similar to Rapamycin, we detected a significant downregulation at mRNA level for CLU, Caspase-3, -8 and -9, p53 and Bax for the ITP plasma treated cells in comparison to control plasma treated cells but a significant increase when compared to plasma untreated cells (Figure 5).

**Figure 5.**
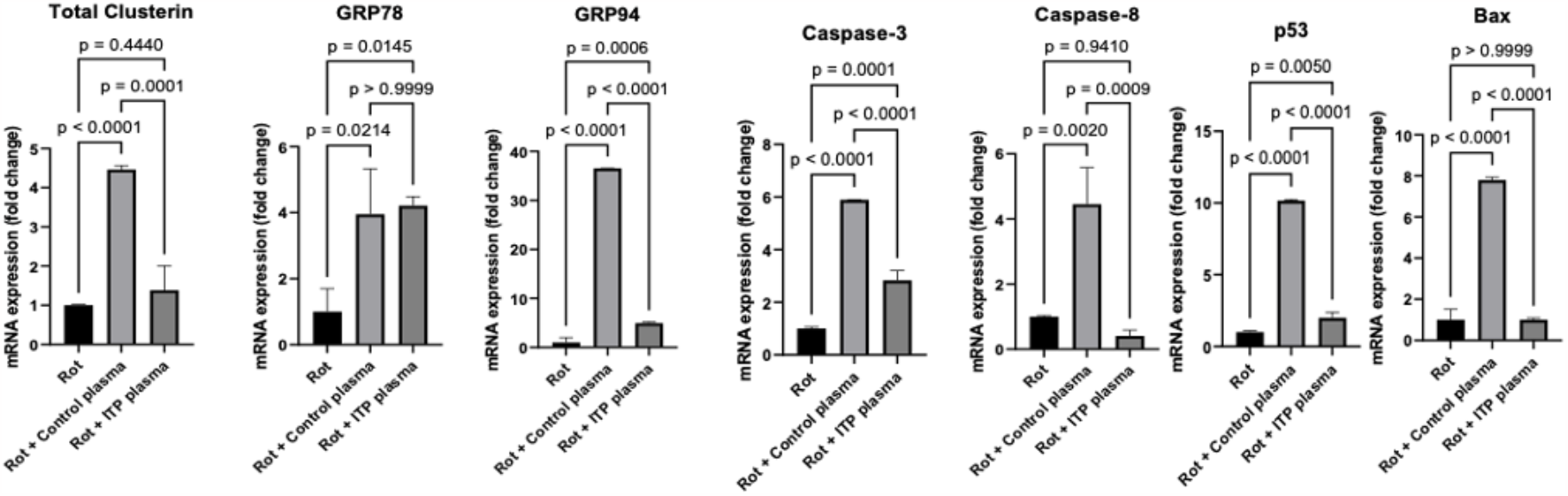
Effects of Rotenone on apoptosis genes in MEG-01 cell line. Apoptosis genes were significantly downregulated in ITP plasma and rotenone treatment. Relative mRNA levels of each sample normalized to control condition are shown as means ± SD. Error bars show SD. Statistical analyses were performed using one-way ANOVA followed by multiple comparisons tests to compare the mean ranks between the groups.

Next, in order to elucidate if the apoptotic death could be caspase-mediated, we performed suppressing apoptosis experiments by treating with pan-caspase inhibitors (Z-VAD-FMK, Promega) and inducing apoptosis experiments by treating with ABT-737, an inhibitor of Bcl-2 family proteins by preventing the sequestration of proapoptotic molecules. In the non- and treated cells we quantified the transcription levels of apoptotic regulatory markers (Figure 6 and 7). As expected, ITP plasma treated cells exhibited an upregulation of all apoptosis markers expression upon ABT-737 treatment which could be reversed when using the pan-caspase inhibitors. These results indicate that the apoptosis-induced mechanisms in ITP could be caspase and Bcl-2 dependent.

**Figure 6.**
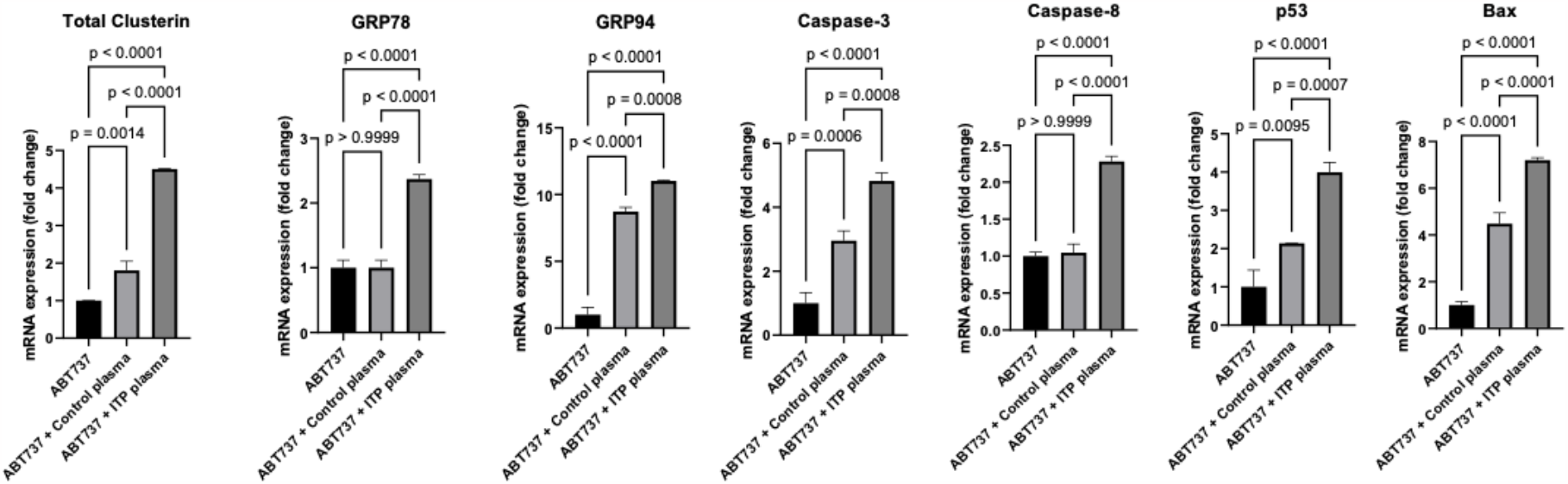
Effects of ABT737 on apoptosis genes in MEG-01 cell line. Apoptosis genes were significantly upregulated in ITP plasma and ABT737 treatment. Relative mRNA levels of each sample normalized to control condition are shown as means ± SD. Error bars show SD. Statistical analyses were performed using one-way ANOVA followed by multiple comparisons tests to compare the mean ranks between the groups.

**Figure 7.**
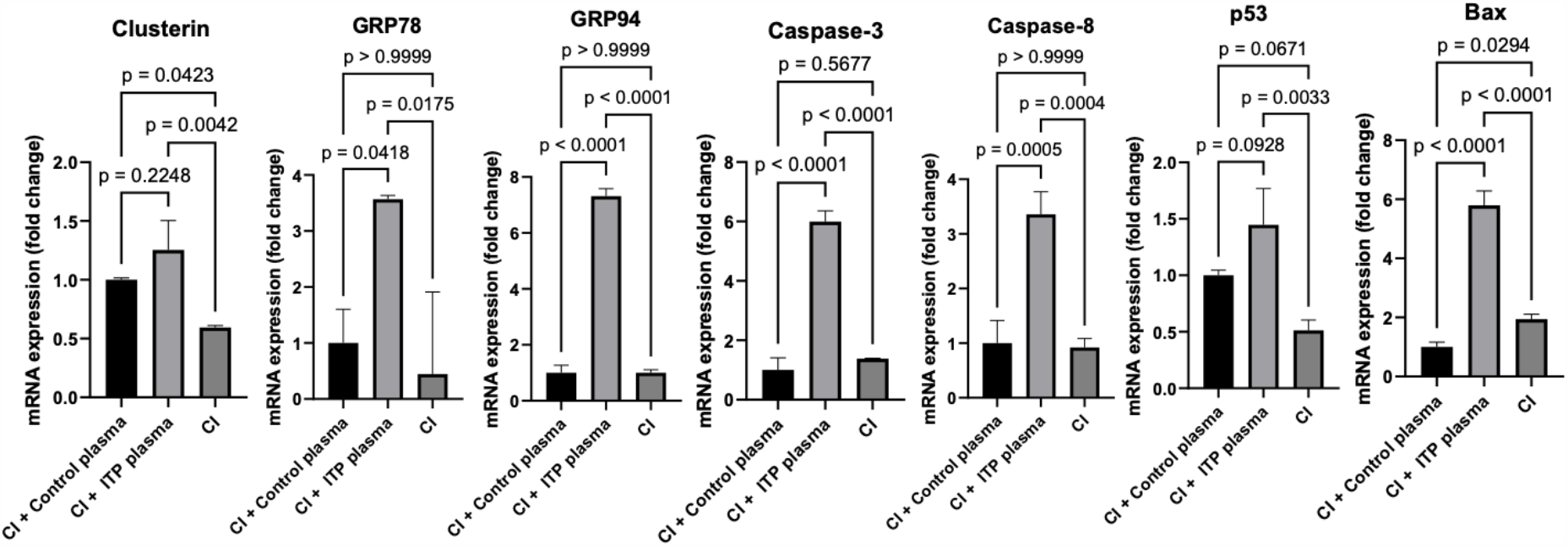
Effects of pan-caspase inhibitor on apoptosis genes in MEG-01 cell line. Apoptosis genes were significantly upregulated in ITP plasma and caspase inhibitors treatment. Relative mRNA levels of each sample normalized to control condition are shown as means ± SD. Error bars show SD. Statistical analyses were performed using one-way ANOVA followed by multiple comparisons tests to compare the mean ranks between the groups.

### CLU siRNA knockdown in ITP plasma treated megakaryocytes

Next, we investigated the CLU function in apoptosis by blocking its expression via RNA interference using CLU siRNA in ITP plasma treated MEG-01 cells. The transfected cells were tested for the expression of CLU, GRP78 and Bax, showing a significant downregulation after ITP plasma treatment compared to untreated or control plasma treatment (Figure 8).

**Figure 8.**
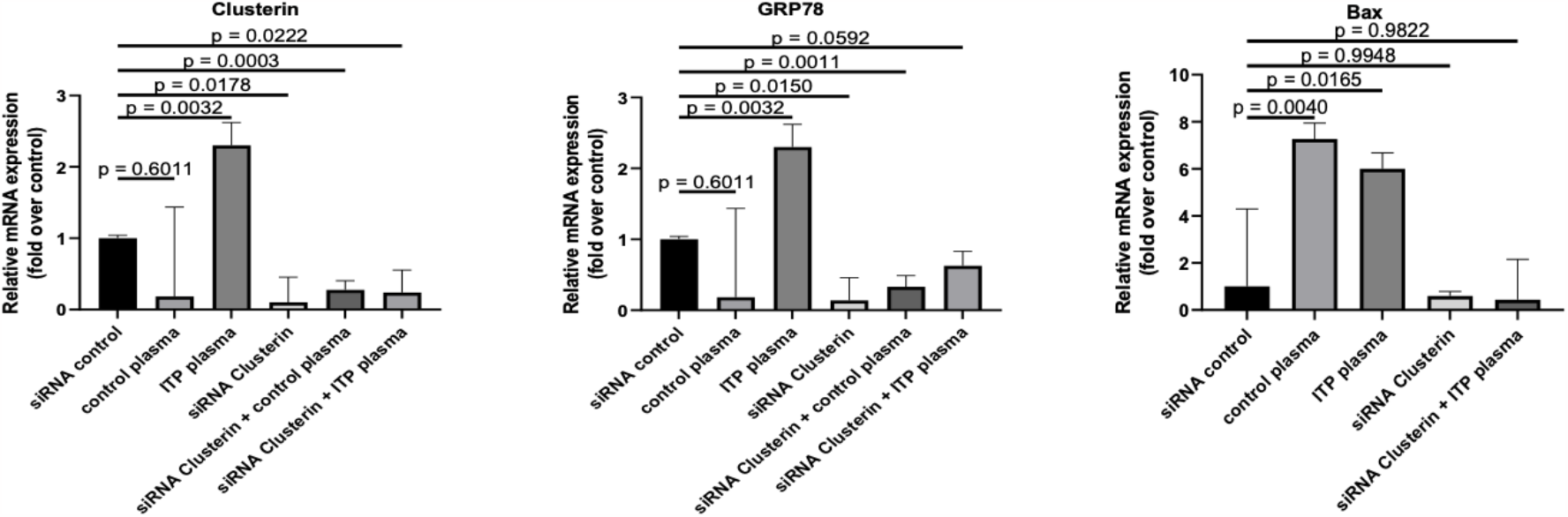
MK cell line MEG-01 treated with ITP plasma shows the different clusterin\ expression after siRNA transfections. mRNA levels of Clusterin determined by qRT-PCR in MEG-01 cell line treated with plasma from ITP patients compared to plasma from healthy Controls and trasfected with siRNA Clusterin. Statistical analyses were performed using one-way ANOVA followed by multiple comparisons tests to compare the mean ranks between groups. Expression levels are shown as means; error bars show SDs and P values are indicated.

## Discussion

ITP is a highly complex autoimmune disease, and its pathogenic mechanisms are still poorly understood. Our first proteomic data revealed that CLU together with other apoptotic markers is significantly upregulated in platelets from both ND and chronic ITP patients compared to healthy controls. To better understand the role of CLU in the pathogenesis of ITP, we investigated possible molecular mechanisms that could lead to CLU activation linked to apoptosis pathway by using different apoptosis inhibitors or inducers as well as siRNA knockdown targeting CLU.

The role of platelet apoptosis into the pathogenesis of ITP has been investigated in some previous research studies with contradictory or inconsistent results in either platelets or MKs, reporting that different regulators such as Akt signaling, Bcl-xL and Bax-mediated apoptosis or PI3K/AKT/mTOR pathway could play a role in ITP^5,30–35^. Our past published work from our research group reported the activation of caspases and phosphatidylserine exposure in ITP platelets^24,36–38^ but the exact mechanisms are still unknown. We could show that ITP plasma treatment on MEG-01 cells could have activating effects on some apoptotic markers including CLU that could be reversed by using either chemical inhibitors or siRNA transfection. Another previous study with plasma treatment, showed decreased expression of Bcl-xL and increased expression of Bax and activity of Caspase-3 in ITP^32^. Deng et al^39^ indicated a significant increase in both surface-exposed PS in platelets and in the expression levels of Bak and Bax in chronic ITP patients compared to healthy controls. Furthermore, Bcl-xL expression levels were significantly decreased in platelets of chronic ITP patients. These results suggested an increased activation of the apoptosis pathway in platelets of chronic ITP patients, which is consistent with our results. Furthermore, it has been shown that ITP plasma is not only involved in platelet destruction but may also play a role in the inhibition of platelet production^40^.

Clusterin (CLU) has been found to be overexpressed in different disease pathologies and used as a therapeutic target Alzheimer’s disease^41,42^, cancer (breast, ovarian and prostate cancer reviewed by^13,42^) or cardiovascular disease (^43^reported as potential prognosis marker for heart damage). In the past years CLU has been targeted as a valid therapy option in these diseases, e.g. using specific drug inhibitors of CLU such as Custirsen OGX-011, a second-generation antisense nucleotide in the treatment of non-small lung cancer, renal cell carcinoma, breast and prostate cancer^44–47^. Recent studies reported the dual function of the CLU isoforms: a proapoptotic nuclear form (nCLU) and a prosurvival secretory form (sCLU)^48^. CLU have been implicated in other diseases, mechanisms where Bax-mediated apoptosis^49^, PI3K-Akt and GSK-3beta signaling^17^ or oxidative stress^50,51^ have been reported as specific pathways that are affected by dysregulated CLU (as reviewed by^42^).

We could validate the proteomic results for CLU at both mRNA and protein level for platelets in ITP samples compared to healthy controls. We could only determine the upregulation of CLU only in ITP platelets and not in PBMCs, which suggests that CLU could exert an apoptotic response only in platelets. CLU has been previously reported to be detected predominately in human platelets, also by immunohistochemistry^20,52^. As CLU mRNA was detected by in situ hybridization in MKs from bone marrow, Tschopp et al^20^ concluded that most probably CLU is produced already during MK development. In line with this, we could detect a similar increase to platelets of mRNA levels of CLU also in MKs treated with ITP plasma compared to control plasma. Mechanistically, GRP78 and CLU are known to directly interact under ER stress conditions, facilitating its redistribution to the mitochondria and trafiking^21,22^. We could also show an increase of the two ER luminal chaperons GRP78 and GRP94 at mRNA level in MKs treated with ITP plasma. Moreover, we could downregulate their expression up to the control levels when using Rapamycin or siRNA targeting either CLU or GRP78. This result could shed more light into the possible mechanism underlying CLU increase in ITP platelets, suggesting that ER-stress induced apoptosis might play a role, as previous reports confirmed their direct interaction also by confocal microscopy and co-immunoprecipitation ^21^.

An interesting study by Zhang et al^53^ revealed that CLU can inhibit apoptosis by interfering with activation of Bax in mitochondria, specifically, CLU interacts with conformation-altered Bax upon treatment with chemotherapeutic drugs. In agreement with this, our experiments showed the same response pattern of CLU and Bax in MEG-01 cells treated with ITP plasma and apoptosis inducers Rapamycin or ABT-737 suggesting that the elevated levels of CLU could interfere with Bax pro-apoptotic activity. Taken together, it will be of interest to elucidate whether disrupting the CLU-Bax interaction may be a possible strategy not only in cancer therapy but as well in ITP disease or other autoimmune disorders.

Rapamycin, also named sirolimus, specifically targets mTOR, a conserved serine-threonine kinase member of the PI3K kinase family regulating cell survival, proliferation and differentiation^54^. It has been reported as treatment in primary and secondary ITP as well as in other autoimmune disorders such as autoimmune lymphoproliferative syndrome (ALPS), systemic lupus erythematosus (SLE) or primary antiphospholipid syndrome (APS)^54^. As favorable clinical evidence regarding Rapamycin treatment in ITP is beginning to accumulate, we were interested to explore its effects on key apoptotic genes expression including CLU when the MK cell line MEG-01 was treated with ITP plasma. The downregulation at mRNA level of all investigated apoptosis genes including Caspase-3, -8, Bax, p53, CLU, GRP78 upon ITP plasma and Rapamycin versus their upregulation after ABT-737 or Rotenone treatment might suggest that Rapamycin could have an anti-apoptotic effect in ITP. This result could be supported by the positive effects of Rapamycin in the clinic, however is in contrast with other reports in retinoblastoma cells^55^, rhabdomyosarcoma cell lines^56^ or in childhood acute lymphoblastic leukemia cells^57^ where Rapamycin was reported to induce apoptosis. Future studies targeting the PI3K/Akt signaling, or p53-dependent mechanisms will shed more light on the impact of these apoptogenic stimuli on MK or platelet survival.

In conclusion, our findings presented here strongly suggest that CLU and its associated molecules GRP78 and GRP94 might have a role in the pathogenesis of ITP, an autoimmune disease. This study contributes to a better understanding of possible apoptosis mechanisms underlying ITP pathogenesis identifying promising molecular targets with potential therapeutic potential.

## Contributions

The study was conceived and coordinated by FDF and MS. The experiments were designed and conducted by TS, CB and FDF. Data analysis was conducted by TS, CB and FDF. The project was supervised by FDF and MS. The manuscript was drafted and refined by TS and FDF with contributions from all authors.

## Acknowledgements

We would like to thank Nadine Goelz, Rosa Hinselmann, Claudia Gowin and David Graber for technical support. This study was founded by Vontobel Foundation and Jacques & Gloria Gossweiler Foundation grants (FDF).

