## Supplementary Figure 1 for "Clusterin can mediate apoptosis-induced molecular mechanisms in immune thrombocytopenia"

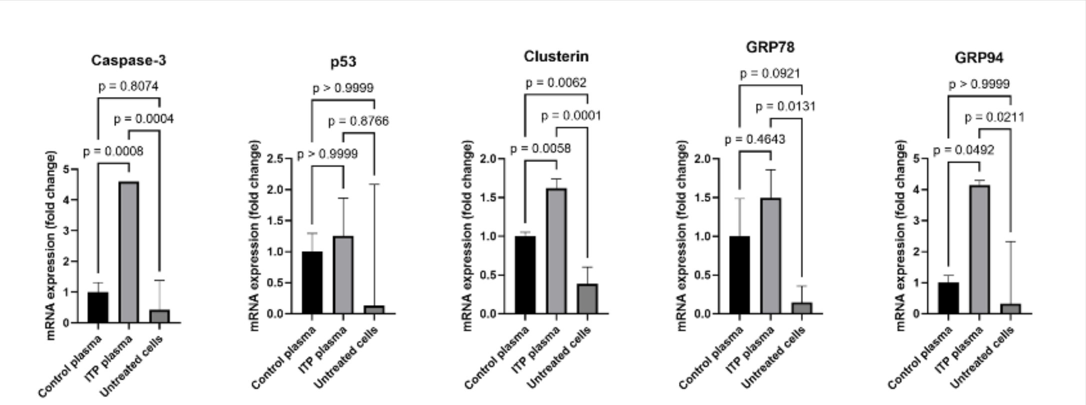


Supplementary Figure 1. Effects of ITP versus control plasma treatment on apoptosis genes in MEG-01 cell line. Apoptosis genes were significantly upregulated in ITP plasma. Relative mRNA levels of each sample normalized to control condition are shown as means ± SD. Error bars show SD. Statistical analyses were performed using one-way ANOVA followed by multiple comparisons tests to compare the mean ranks between the groups.
